# Revising transcriptome assemblies with phylogenetic information in Agalma1.0

**DOI:** 10.1101/202416

**Authors:** August Guang, Mark Howison, Felipe Zapata, Charles Lawrence, Casey Dunn

**Affiliations:** Division of Applied Mathematics, Brown University, Providence, RI 02912, USA; Computing & Information Services, Brown University, Providence, RI 02912, USA; Department of Ecology & Evolutionary Biology, University of California-Los Angeles, Los Angeles, CA 90095, USA; Department of Ecology & Evolutionary Biology, Yale University, New Haven, CT 06511, USA.

## Abstract

**Motivation:** One of the most common transcriptome assembly errors is to mistake different transcripts of the same gene as transcripts from multiple closely related genes. It is difficult to identify these errors during assembly, but in a phylogenetic analysis these errors can be diagnosed from gene trees containing clades of tips from the same species with improbably short branch lengths.

**Results:** `treeinform` is a module implemented in Agalma1.0 that uses phylogenetic analyses across species to refine transcriptome assemblies. It identifies transcripts of the same gene that were incorrectly assigned to multiple genes and reassign them as transcripts of the same gene.

**Availability and Implementation:** `treeinform` is implemented in Agalma1.0, available at https://bitbucket.org/caseywdunn/agalma.

**Supplementary information:** Supplementary information is available at bioRxiv.

## 1 Introduction

The development of RNA-seq has made it possible to generate large amounts of transcriptome data for a broad diversity of species, providing novel insights into the history of organisms and the study of gene evolution and function (Wang *et al.*, 2009). The central goal of transcriptome assembly is to use the raw reads to estimate the correct sequence of the original transcripts and assign these to their respective genes. One of the biggest challenges in transcriptome assembly is to identify whether sequence variance is due to technical factors (such as errors introduced in library prep and sequencing), splicing differences, different alleles of the same locus, or evolutionary divergence between closely related gene copies following duplication. This makes it difficult to determine if slightly different transcripts are variants of the same gene or are derived from different closely related genes. We assessed the prevalence of this problem in a test dataset and found that it is extremely common (see Supplementary Figure 1). When different transcripts of the same gene are incorrectly assigned to multiple genes, downstream analyses (e.g., phylogenetics, gene expression) can be compromised.

When transcriptomes are compared across species in a phylogenetic framework, transcripts of the same gene that have been incorrectly assigned to multiple genes will appear in gene trees as tips with improbably short branch lengths (Figure 1). Here we present a method, `treeinform`, that uses this phylogenetic information to identify misassigned transcripts and reassign them to the same gene. Our approach uses a subtree length threshold and revises the mapping of transcripts to genes accordingly. `treeinform` essentially borrows information across species to refine the assignment of transcripts to genes within species.

**Figure 1:**
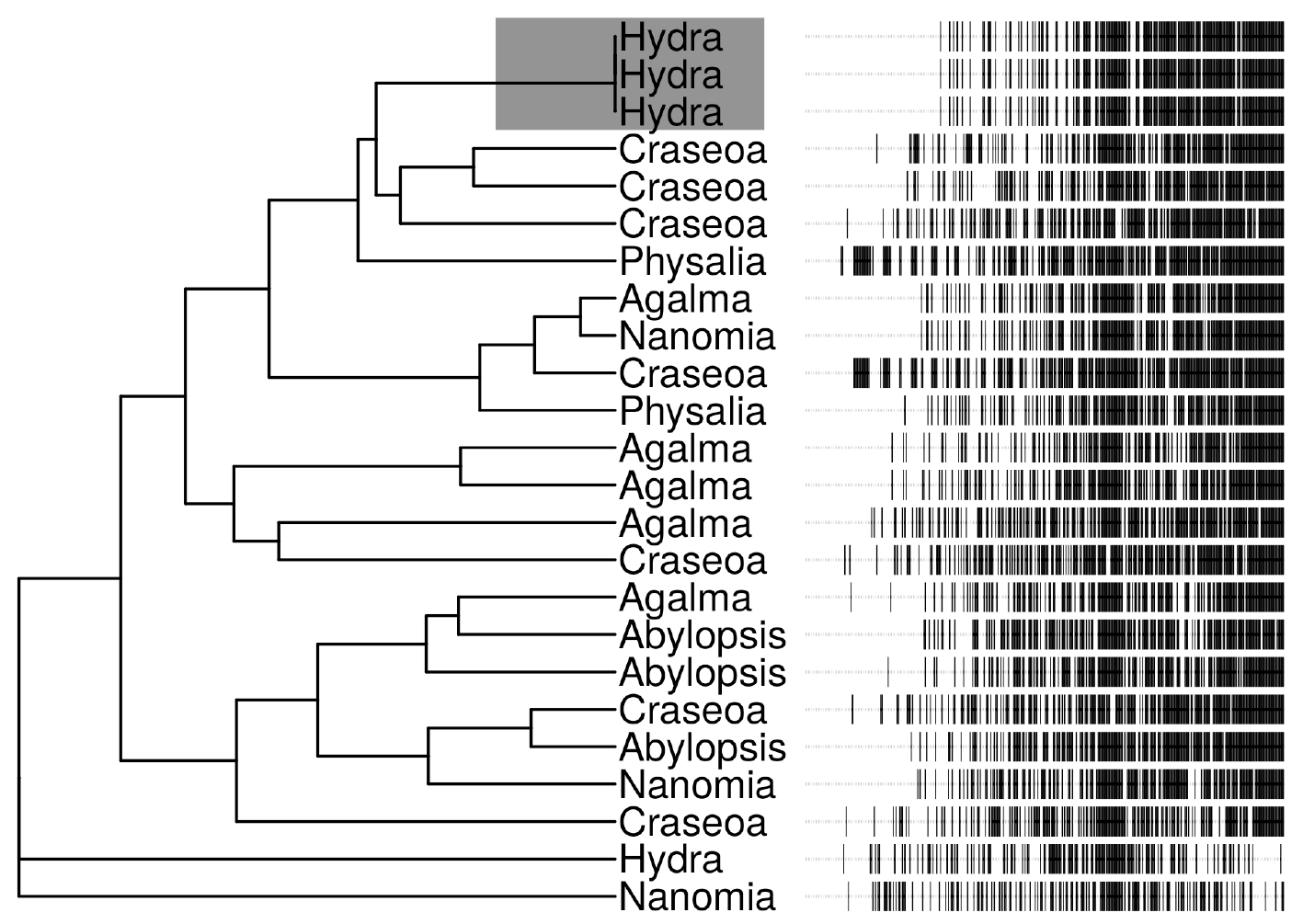
On the left is a gene tree from the test dataset before running `treeinform`. Each tip is an exemplar transcript that was initially assigned to a different gene. On the right is the multiple sequence alignment, with sequence sites ordered from highest to lowest identity to the inferred ancestral site for clarity on sequence diversity. Black indicates a difference from the ancestral sequence. The three *Hydra* transcripts in the grey box at the top of the gene tree were assigned to different genes by Trinity (Grabherr *et al.*, 2011) despite having nearly indistinguishable sequences. After `treeinform`, all transcripts from these three genes would be reassigned to a single gene.

## 2 treeinform Overview

### 2.1 Method

`treeinform` is implemented as a module within the phylogenomic workflow Agalma1.0 (Dunn *et al.*, 2013), which includes a variety of other updates such as increased portability of analyses and greatly simplified installation. For each internal node on a tree, subtree length is calculated as the sum of all branch lengths in the subtree defined by the node. `treeinform` then identifies subtrees with total length below a given threshold. If multiple tips (i.e., genes) belonging to a single species exist in an identified subtree, all transcripts for these multiple genes are flagged for reassignment to a single gene. All reassigned transcripts are rerun through the pipelines in Agalma to complete a phylogenomic analysis.

### 2.2 Input/Output

Input consists of a set of gene trees generated by the pipeline `genetree` in Agalma1.0 and a user-designated threshold. The default value for the threshold is 0.02, selected based on analyses of a set of species for which the species tree is well-resolved (see Supplementary Information). The user may want to run their own analyses to determine an appropriate threshold. `treeinform` outputs a list of transcripts for reassignment, which become input for a second run of Agalma1.0 starting from RSEMEval (Li *et al.*, 2014).

### 2.3 Results

We tested `treeinform` on a dataset of seven species of siphonophores. We took two different approaches to assess the efficacy of `treeinform`. First, we spot checked the results to confirm that they were biologically and technically sensible. This provided detailed confirmation on a small fraction of the output. Second, we compared duplication time distributions of the entire input and output tree sets against theoretical expectations (Gernhard, 2008) (see Supplementary Figure 3). Duplication times are related to subtree lengths because when transcripts from the same gene are misassigned to different genes, gene tree/species tree reconciliation programs compensate by inferring additional duplication events. Because such transcripts have almost identical sequences, the inferred duplication events will be extremely shallow, with correspondingly recent duplication times. Conversely, correctly assigned transcripts from different genes will have less similar sequences and thus older duplication times. Duplication time distributions of output tree sets under the default threshold were much closer to theoretical expectations than from input tree sets (see Supplementary Information). This provided an overview of the impact of the method across the entire output.

## 3 Conclusion

We confirm that assignment of transcripts from the same gene to different genes is quite common in transcriptome assemblies. Using phylogenetic information, our new approach reassigns transcripts to their corresponding gene when different transcripts of the same gene are mistaken as transcripts from different closely related genes. This approach capitalizes on the Markovian dependency assumption in phylogenetic workflows (Guang *et al.*, 2016), as not only can inferences about the gene trees be made solely from the gene assemblies, but inferences about the gene assemblies can be made solely from the gene trees as well. Analyses of `treeinform` shows that it brings estimates of duplication times much closer to theoretical expectations. `treeinform` will be useful for any studies requiring accurate gene trees, in particular accurate counts of different genes in expression studies. It is available in Agalma1.0, an end-to-end phylogenomic workflow.

## Acknowledgements

Thanks to Alejandro Damian Serrano for feedback on the manuscript.

